# A dynamically coupled kinetic model for Arterial Spin Labeling: Enhancing the reliability of ventricular cerebrospinal fluid renewal quantification in vivo

**DOI:** 10.64898/2026.07.28.741371

**Authors:** Qiong Ye

## Abstract

Arterial spin labeling (ASL) provides a valuable non-invasive tool for investigating cerebrospinal fluid (CSF) dynamics. While existing generalized kinetic models provide a useful analytical framework, incorporating explicit fluid mass-conservation constraints may improve the reliability of ventricular CSF quantification. We developed and validated a physics-constrained kinetic model that assumes a constant ventricular volume, under which the volumetric influx and efflux rates are balanced. Under this assumption, the localized CSF renewal rate (*f*’) is modeled as a distinct washout process that acts jointly with intrinsic CSF T_1_ relaxation *R*_1,*CSF*_, yielding an effective decay rate *R*_*eff*_ = *R*_1,*CSF*_ + *f*’, providing a more mechanistically interpretable description of the post-arrival signal decay. The model was further tailored to the global inversion physics of the FAIR (Flow-sensitive Alternating Inversion Recovery) sequence and incorporates an explicit zero-clamped pre-arrival boundary condition. When applied to an in vivo preclinical dataset, the proposed model, improved fitting stability and removed the finite-bolus truncation observed in the raw conventional model. Compared with the conventional model, the proposed model showed significantly improved goodness-of-fit (*R*^2^ = 0.95 ± 0.05 vs. 0.82 ± 0.06, *p* < 0.001) and a lower Akaike Information Criterion (*AIC* = 94.96 ± 5.83 vs. 106.93 ± 2.50, *p* < 0.001), suggesting it provides a more adequate and efficient representation of CSF dynamics. The proposed model yielded CSF dynamics estimates 14.70% higher than those obtained with the conventional model, with a mean ventricular CSF renewal time of 4.01 ± 0.97 min and a renewal-equivalent volumetric flow rate of 0.97 ± 0.41 *µ*L/min. The resulting estimates might be interpreted as localized, ASL-derived renewal metrics rather than direct measurements of net CSF production.

## Introduction

The efficient clearance of metabolic waste from the central nervous system is critical for maintaining cerebral homeostasis[1-3]. Recently, non-invasive assessment of cerebrospinal fluid (CSF) dynamics using Arterial Spin Labeling (ASL) has emerged as a powerful approach to investigate blood-to-CSF water transport mechanisms[4-9]. By using ultra-long echo time (TE) acquisitions to suppress vascular and parenchymal background signals, the ASL signal associated with water transport across the blood-CSF barrier could be selectively isolated [4, 6]. This technique is named as blood-CSF barrier ASL (BCSFB-ASL). It enables quantitative assessment of blood-to-CSF water transport dynamics within the ventricular system.

Emerging studies have demonstrated the promising value of this technique across both rodent models and human cohorts[4, 6, 8, 9]. Notably, BCSFB-ASL has shown remarkable sensitivity to age-related pathological changes. In mice, the rate of blood-to-CSF water transport declines significantly with aging, a reduction that is often more pronounced than the subtle decreases observed in cortical perfusion, and which occurs prior to any macroscopic ventricular enlargement[4]. Furthermore, in transgenic mouse model of Alzheimer’s disease, BCSFB dysfunction detected by ASL precedes behavioral deficits and amyloid pathology, highlighting it as a crucial early event in neurodegeneration[8]. Parallel investigation in humans has yielded similarly encouraging results[6], collectively suggesting that this technique has considerable potential as an early imaging biomarker for multiple neurological disorders.

Despite these promising applications, the physiological interpretation of ASL-derived metrics requires rigorous clarification. A common misconception is to equate the measured blood-to-CSF water transport with macroscopic net CSF production. In reality, previous studies have confirmed that, in addition to blood-to-CSF water transport, blood-CSF water exchange is a rapid, continuous, and bidirectional process[10]. BCSFB-ASL inherently captures both the transport and exchange of water, rather than net volumetric secretion [1]. Consequently, although invasive methods report very low net lateral ventricular CSF production[11], the ASL-derived effective water flux may be orders of magnitude larger because it reflects both water transport and rapid bidirectional water exchange rather than net volumetric secretion[4]. To avoid semantic ambiguity with the conventional concept of global volumetric CSF turnover and to mitigate the risk of unwarranted physiological extrapolation, this ASL-derived metric is more appropriately defined as the **localized CSF renewal rate (***f*′), a measure reflecting the dynamic fractional replacement of fluid within a specific anatomical compartment [1].

Despite its physiological interpretability, the widespread adoption of BCSFB-ASL remains hindered by several technical bottlenecks. Inherent limitations include a low signal-to-noise ratio (SNR), spatial resolution constraints, and the prolonged scan times required for multi-delay acquisitions. Beyond these challenges, the quantification of ASL-derived metrics is highly sensitive to the underlying kinetic model used. Existing kinetic models often do not explicitly account for coupled influx-efflux dynamics under fixed-volume condition, nor do they fully reflect sequence-specific boundary condition, which can increase parameter uncertainty [4, 8]. Consequently, these modeling vulnerabilities might cause inter-study variability, ultimately undermining the reliability of ASL as a quantitative measure of CSF dynamics. Therefore, a refined analytical model grounded in fluid mass conservation and actual sequence physics is essential to improve the robustness and physiological relevance of the methodology.

To address these critical methodological challenges, we propose a novel, physiologically constrained analytical model: a dynamically coupled kinetic model for ASL-based quantification of CSF renewal rate within the lateral ventricles. Our model is specifically tailored to the Flow-sensitive Alternating Inversion Recovery (FAIR) ASL sequence, which is widely adopted in preclinical imaging[12, 13]. Compared with the conventional three-parameter model, our approach introduces two fundamental refinements. First, governed by the principle of fluid mass conservation, we dynamically couple fluid influx and efflux, thereby explicitly isolating the physiological renewal rate from the intrinsic *T*_1_ relaxation of the CSF. Second, by anchoring the model to the underlying physics of the global inversion sequence, we formulate the unlabeled water input as a continuous steady-state process (*τ* → ∞), thereby naturally reducing the model to a more stable two-parameter space (renewal rate *f*’ and transit time *δ*).

Here, we systematically evaluated this refined model against the conventional model using Monte Carlo simulations and an open-access in vivo mouse MRI data. Our results demonstrate that the mass-conserved model effectively mitigates quantification bias, eliminates artifactual curve truncations, and improves fitting performance. This yields physiologically more plausible washout profiles and superior goodness-of-fit. By providing a highly robust assessment of localized CSF renewal, this model expands the translational utility of ASL for investigating ventricular CSF dynamics in the context of aging and neurodegenerative disorders.

### Theory

FAIR-ASL measures the difference between two interleaved inversion preparations, typically a slice-selective (label) and a non-selective/global inversion condition (control). In the present implementation, the ASL difference signal was defined as *ΔM* = *M*_*label*_ − *M*_*control*_, such that a positive signal represents the arrival of uninverted blood-derived water into the ventricular CSF compartment. Under ultra-long TE acquisition, vascular and parenchymal signals are strongly attenuated, and the remaining ventricular signal is assumed to primarily reflect blood-derived water transport and exchange with CSF.

As illustrated in Figure 1, we model the lateral ventricles as a well-mixed compartment with constant macroscopic volume (*V*_*LV*_) during the ASL experiment. Under this assumption, the volumetric influx and efflux of CSF are balanced, such that *Q*_*in*_ = *Q*_*out*_ = *Q*. The parameter *f*′(*s*^−1^) represents the localized CSF renewal rate, defined as the fractional replacement rate of ventricular CSF water by blood-derived water per unit lateral ventricular volume; that is, the rate at which pre-existing ventricular CSF water is replaced by newly arrived blood-derived water. Physiologically, this process may involve water transport across the choroid plexus, as well as water exchange across the lateral ventricular wall. The constant-volume assumption should be interpreted as a time-averaged approximation over the ASL observation window, rather than as an absence of pulsatile CSF motion.

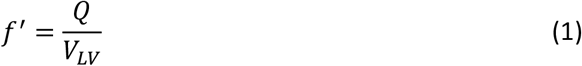

which, under the fixed-volume assumption, is also equal to the efflux-driven washout constant,

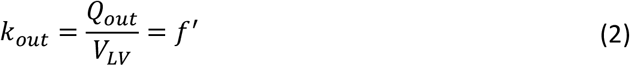

**Figure 1.**
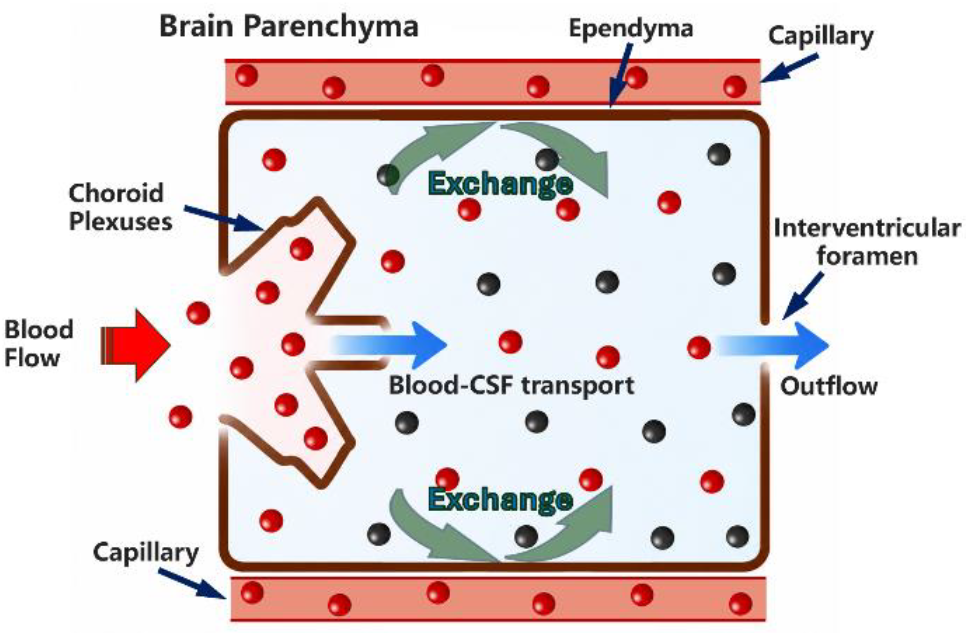
Schematic diagram of the ventricular CSF water renewal model. The lateral ventricle is treated as a well-mixed compartment with a constant volume. Black spheres denote pre-existing CSF water, and red spheres denote newly arrived blood-derived water. Water enters the ventricle via blood-CSF transport across the choroid plexus and exchange across the ependymal/lateral ventricular wall, while compensatory bulk outflow through the interventricular foramen maintains a constant ventricular volume.

Under this condition, the kinetic equation for the label-control difference signal *ΔM*(*t*) of the CSF in the lateral ventricles is

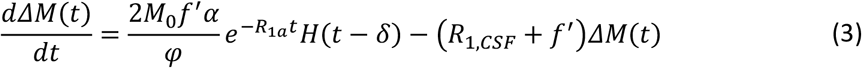

Here, *M*_0_ is the equilibrium magnetization of ventricular CSF and *R*_1,*CSF*_ is its longitudinal relaxation rate. *M*_0_ and *R*_1,*CSF*_ were estimated independently from the control signal. In addition, *α* is the labeling efficiency, *φ* is the blood-CSF partition coefficient (assumed to be 1), *R*_1*a*_ is the longitudinal relaxation rate of blood, *H*(*t* − *δ*) is the Heaviside step function, and *δ* is the transit delay for unlabeled blood water to reach the CSF in the lateral ventricles.

In Eq. (3), time *t* denotes the time elapsed since the inversion pulse. The factor 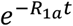 accounts for the longitudinal relaxation of the inflowing blood-derived water from the time of inversion to the time at which it contributes to the ventricular CSF signal. The Heaviside function *H*(*t* − *δ*) imposes the transit delay, such that the input term becomes active only after the arrival of uninverted water into the ventricular compartment. Notably, the model distinguishes between conservation of fluid volume and washout of difference magnetization: constant ventricular volume implies equal volumetric influx and efflux, whereas the efflux term for the difference signal follows from a well-mixed compartment assumption, under which the exiting fluid carries the mean compartmental difference magnetization. The ventricular difference magnetization therefore follows a tracer-balance form and decays through intrinsic CSF longitudinal relaxation and replacement-driven washout, giving the combined decay constant *R*_1,*CSF*_ + *f*′.

The localized CSF renewal time (LCRT) is defined as

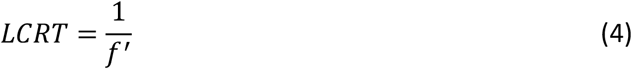

Given the boundary condition *ΔM*(*δ*) = 0, the exact solution to the model is:

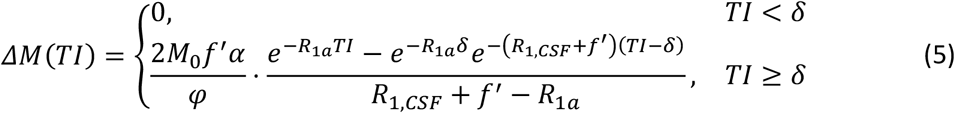

The complete analytical derivation is given in the Supplementary Material.

For comparison, the conventional generalized kinetic model described by Evans et al. [4] is expressed as

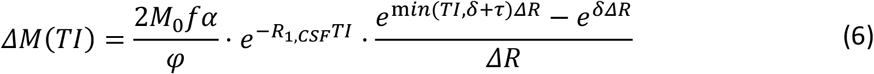

Where

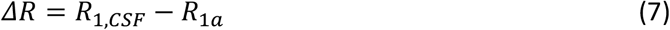

and *τ* is the label duration, as employed in PASL and pCASL.

Based on the literature [4], *f* denotes the rate of delivery of labelled blood water to ventricular CSF and serves as a quantitative surrogate marker of BCSFB function, primarily reflecting blood-to-CSF water transport. Although in the original study *f* reflected only blood-to-CSF water transport, the ventricular CSF signal intensity is also affected by water exchange between the CSF in the lateral ventricle and the lateral ventricular wall. Therefore, *f* reflects both water transport and exchange, which is consistent with *f*′. In its original implementation, this expression does not explicitly enforce *ΔM* = 0 for *TI* < *δ*, and the finite bolus duration produces a change in slope at *TI* = *δ* + *τ*. These features were retained here for direct comparison with the raw generalized model.

## Materials and methods

### 2.1 Kinetic models and numerical simulations

To systematically examine parameter recovery within the proposed model and to illustrate the artificial curve truncation that may arise when the data are fitted with the conventional generalized model [4], numerical simulations were performed using MATLAB (MathWorks, Natick, MA). Prior to simulation, our proposed dynamically coupled two-parameter model was formulated upon the following strict physiological and sequence-specific assumptions:

1. Constant compartmental volume: The macroscopic volume of the lateral ventricles remains constant throughout the ASL measurement.
2. Balanced mass flow: Under the fixed-volume assumption, the volumetric rate of blood-derived water entering the compartment through transport and exchange is balanced by the volumetric rate leaving the compartment.
3. Distinct but coupled loss processes: The physiological washout term *f*′ is modeled explicitly as a process distinct from intrinsic CSF T_1_ relaxation, while both jointly contribute to signal decay.
4. Zero-clamp boundary condition: The model explicitly incorporates the transit phase (*TI* < *δ*), enforcing a physically realistic zero-signal constraint before the arrival of fresh, unlabeled blood.
5. Continuous flow dynamics: Tailored to the global inversion pulse of the FAIR sequence, the labeled bolus duration is assumed to be infinite (*τ* → ∞).

For clarity, the estimated parameters in the conventional and proposed models are defined as follows. In the conventional three-parameter model, the estimated parameters are the transport rate of labelled blood water to ventricular CSF, normalized by ventricular CSF volume (*f*), transit time (*δ*), and label duration (*τ*). In the proposed two-parameter model, the estimated parameters are the localized CSF renewal rate (*f*′), defined as the fractional replacement rate of ventricular CSF by blood-derived water per unit lateral ventricular volume, and transit time (*δ*). Although *f* and *f*′ are defined differently, both are derived from changes in the lateral ventricular water signal, which is assumed to be influenced not only by BCSFB-mediated delivery of blood-derived water but also by water exchange across the lateral ventricular wall and bulk water flow through the ventricular foramina. Therefore, although these processes are not explicitly included in the original three-parameter model formulation, they may all affect parameter estimation. From this perspective, both *f* and *f*′ can be regarded as model-based measures related to CSF dynamics within the lateral ventricles.

#### 2.1.1 Simulation setup and noise-free signal-shape evaluation

Simulations were performed using predefined global parameters for a 9.4 T field strength, including equilibrium magnetization *M*_0_ = 1000, inversion efficiency *α* = 0.9, partition coefficient *φ* = 1.0, arterial longitudinal relaxation time *T*_1*a*_ = 1⁄*R*_1*a*_ = 2.4 s [14], and ventricular CSF longitudinal relaxation time *T*_1,*CSF*_ = 1⁄*R*_1,*CSF*_ = 4.0 s. The ground-truth localized CSF renewal rate and transit delay were set to 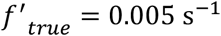 and *δ*_*true*_ = 0.50 s, respectively. For the conventional three-parameter model, the ground-truth values were set to 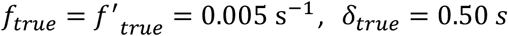 and *τ*_*true*_ = 4.0 *s*, respectively. The temporal sampling schedule matched a typical ultra-long TE FAIR-ASL protocol, consisting of six discrete inversion times (TIs): [0.2, 0.75, 1.5, 2.75, 4.0, 6.5] s.

To evaluate and compare the theoretical differences arising from the two mathematical models, a noise-free signal-shape analysis was first conducted. The noise-free ground-truth signals at the six TIs were synthesized according to the proposed model (Equation 5) and the conventional three-parameter model (Equation 6), respectively.

#### 2.1.2 Monte Carlo noise robustness analysis

To investigate the robustness of the proposed two-parameter model under typical MRI noise conditions, Monte Carlo simulations were performed. Gaussian white noise was added onto the six discrete ground-truth signal points to generate four signal-to-noise ratio (SNR) levels: 15, 30, 60, and 120. Here, SNR was defined as the ratio of the maximum theoretical signal peak to the standard deviation of the added Gaussian noise. For each SNR level, 3000 independent iterations were performed. The proposed coupled two-parameter model was then used to estimate the localized CSF renewal rate (*f*′) from the noisy synthetic data. To avoid non-physiological solutions during optimization, physiologically plausible parameter bounds were imposed during fitting (*f*′ ∈ [0, 0.05]s^−1^ and *δ* ∈ [0, 6.5]s). The distribution of the fitted *f*′ values across the 3000 iterations was visualized using boxplots to assess the accuracy and stability of parameter recovery under noisy conditions.

### 2.2 *In vivo* data acquisition (open-source dataset)

To validate the proposed two-parameter model *in vivo*, we utilized an open-source 9.4T MRI dataset from 12 mice (female C57BL/6J, 3 months), originally described by the study [4]. Guided by T2-weighted anatomical images, a single 2.4-mm coronal ASL slice was centered on the lateral ventricles. CSF dynamics were captured using a FAIR labeling scheme with a wide slice-selective inversion slab (19.2 mm). Importantly, an ultra-long TE of 220 ms was implemented to isolate the CSF signal by effectively suppressing parenchymal and vascular background contributions. ASL data were acquired across six inversion times (TIs = 0.2, 0.75, 1.5, 2.75, 4.0, and 6.5 s) with a repetition time (TR) of 12 s, FOV of 20 × 20 mm^2^, and a 32 × 32 matrix. To compensate for the inherently low SNR at this ultra-long TE, 20 averages were collected per TI. Comprehensive hardware and acquisition details are available in the original study[4]. No new animal experiments were performed in this study. All analyses were conducted using publicly available datasets. Ethical approval for the original animal experiments was obtained by the respective data contributors, as described in the original publication.

### 2.3 *In vivo* kinetic modeling

The 3D volumes of the lateral ventricles were derived from T2-weighted images (T2WI). Binary CSF masks were manually segmented across consecutive slices using ImageJ (NIH, USA), and the 3D volumes were calculated by multiplying the total number of voxels by the voxel volume. For the ASL data, the acquired NIfTl images were initially spatially upsampled by a factor of two, followed by motion correction. In this study, the ASL difference signal was defined as Δ*M*=*M*_*label*_ − *M*_*control*_, where the label image corresponds to the slice-selective inversion condition and the control image corresponds to the non-selective/global inversion condition. To derive ventricular blood-to-CSF signal dynamics, a region of interest (ROl) delineating the lateral ventricles was manually defined within the ASL space. The *ΔM* kinetic time course across the six discrete TIs was then extracted from this ROl for subsequent mathematical modeling. Furthermore, to mitigate partial volume effects (PVE) inherent in low-resolution ASL, a corrected 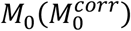 was calculated. This was achieved by scaling the original *M*_0_ signal by the ratio of the high-resolution T2WI-derived ventricular volume to the corresponding ASL-derived ROl volume.

The extracted *in vivo* time-series data were fitted using the proposed dynamically coupled two-parameter model. Optimization was performed utilizing a non-linear least-squares curve fitting algorithm (lsqcurvefit in MATLAB, MathWorks, Natick, MA). For the 9.4T preclinical environment, the longitudinal relaxation time were fixed based on established literature values: 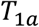 was set to 2.4 s[14]. The blood-brain partition coefficient for water (ϕ) was assumed to be 1.0, and the inversion efficiency (*α*) was not fitted but instead assumed to be constant and fixed at 0.9[4]. The proposed model concurrently estimated two physiological parameters: the localized CSF renewal rate (*f*′) and the arterial transit time (δ).

For comparison, the same *in vivo* datasets were also fitted using the conventional uncoupled three-parameter model (estimating *f*, δ, and the bolus duration *τ*). Consistent with our numerical simulation protocol, the conventional model was executed in its raw mathematical form (Equation 6). The comparative analysis focused on evaluating the fitting performance (Goodness-of-Fit, *R*^2^, and Akaike Information Criterion, AIC) and the physiological plausibility of the resultant kinetic curves.

### 2.4 Statistics

Statistical analyses were performed using SPSS Statistics version 27.0 (IBM Corp., Armonk, NY, USA). Normality was assessed using the Shapiro-Wilk test. Paired two-tailed t-tests were used for normally distributed paired comparisons; otherwise, Wilcoxon signed-rank tests were applied. A *p* < 0.05 was considered statistically significant.

## Results

### 3.1 Noise-free kinetic curves

To elucidate the theoretical distinctions between the two analytical models, we first examined their kinetic signal profiles under noise-free conditions (Figure 2A). The conventional uncoupled three-parameter model has been widely used to characterize ASL kinetics and remains a valuable reference model. In the raw conventional implementation used for comparison, the finite bolus duration introduces a change in kinetic behavior. Although later refinements have partly addressed boundary-condition issues [8], this finite-bolus structure may still be suboptimal for the present FAIR acquisition.

**Figure 2.**
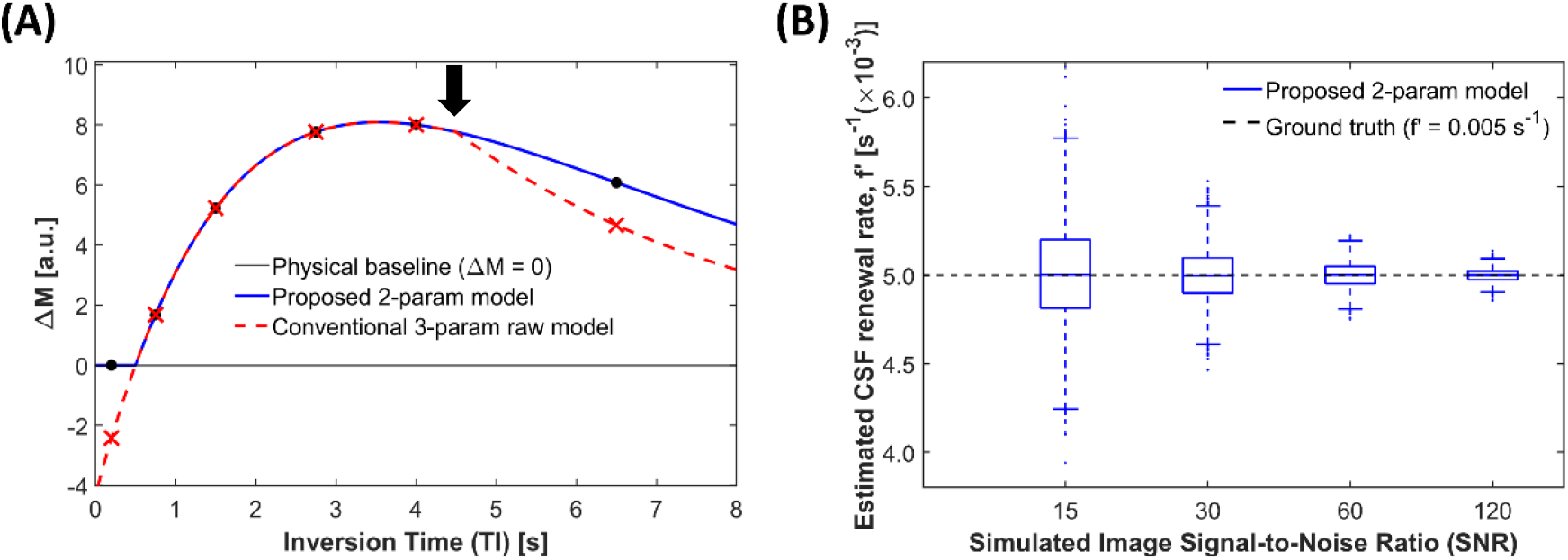
(A) Comparison of noise-free kinetic curves between the proposed two-parameter model (solid blue line, black dots) and the conventional three-parameter model (dashed red line, red crosses). The black arrow indicates that the raw conventional finite-bolus model exhibits an abrupt change in kinetic behavior. (B) Boxplots of the estimated localized CSF renewal rates (*f*′) derived from Monte Carlo simulations across varying SNR levels (15, 30, 60, and 120). The dashed black line indicates the ground-truth value 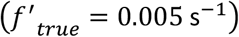.

By comparison, the proposed dynamically coupled two-parameter model is designed to provide a more physiologically consistent model for the present application. Specifically, it incorporates volume conservation within the lateral ventricular compartment and is derived under the limiting condition of *τ* → ∞, thereby eliminating the need to impose an artificial finite bolus termination. This leads to a smoother kinetic profile, with signal evolution determined continuously by influx, T_1_ relaxation, and efflux. Consequently, the proposed model is expected to provide a more natural description of CSF dynamics in the presence of these mechanism, while retaining the conceptual strengths of the established model.

### 3.2 Robustness and parameter identifiability under noise

Monte Carlo simulations under typical MRI noise conditions (Figure 2B) showed that the proposed two-parameter model yielded estimates of the localized CSF renewal rate (*f*′) that were consistently centered near the ground-truth (0.005 s^−1^) across SNR levels of 15, 30, 60, and 120, with decreasing variance as SNR increased.

### 3.3 *In vivo* quantification of CSF dynamics in the lateral ventricles

The proposed model was applied to the open-source in vivo preclinical dataset reported by Evans et al. [4] (Figure 3). Following data quality review, one subject was excluded due to an artifact at the first Tl, leaving a cohort of *N* = 11 for quantitative analysis

**Figure 3.**
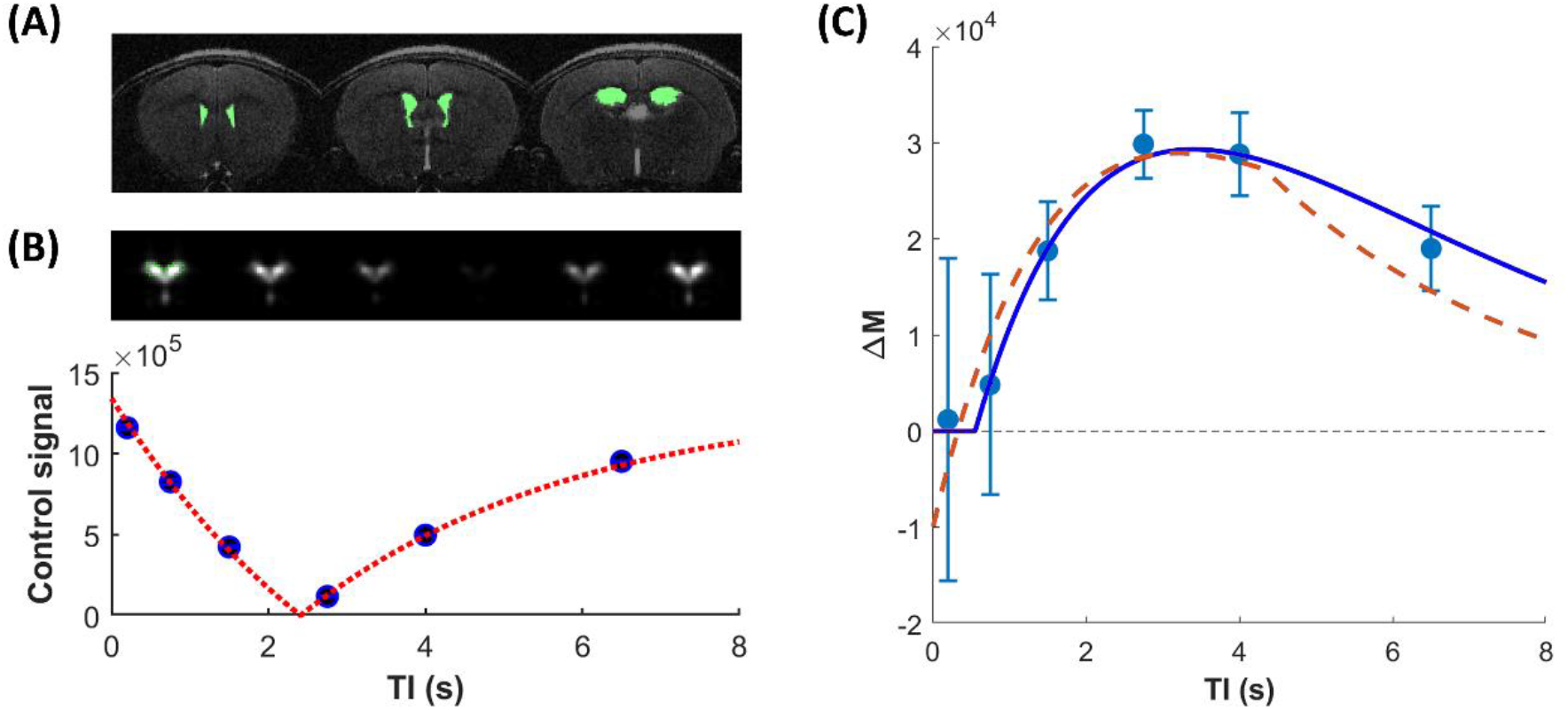
(A) Representative T2-weighted images with the lateral ventricles highlighted in green. (B) Representative control signals across inversion times (TIs) and the corresponding fitted curve. The lateral ventricular ROI is outlined with a green line. (C) ΔM signals of the lateral ventricular ROI across TIs (light blue dots), fitted with the proposed two-parameter model (blue solid line) and the conventional three-parameter model (red dashed line).

Representative T2-weighted images with the lateral ventricles delineated in green are shown in Figure 3A. Across the cohort, the mean lateral ventricular volume was 3.57 ± 0.87*µ*L. The longitudinal relaxation curve, fitted from the control signals across varying Tls (Figure 3B), yielded a mean ventricular CSF *T*_1,*CSF*_ of 3.51 ± 0.24 *s*. Figure 3C illustrates the representative kinetic fitting of the *ΔM* signals using both analytical models. All fitted curves using the proposed two-parameter model are provided in the Supplementary Materials. When applied to the in vivo ultra-long TE ASL data, the conventional three-parameter model estimated the CSF dynamic metric *f* of (3.81 ± 0.92) × 10^−3^ *s*^−1^ and an arterial transit time (*δ*) of 0.28 ± 0.07 s. In contrast, the proposed two-parameter model yielded a *δ* of 0.55 ± 0.08 s and a significantly higher CSF dynamic metric *f*′ of (4.37 ± 1.02) × 10^−3^ *s*^−1^(*p* < 0.001), representing a 14.70% increase in the estimated CSF dynamic metric. Based on the proposed model, the mean localized CSF renewal time was 4.01 ± 0.97 minutes. By scaling *f*′ with individual ventricular volumes, this translated to an absolute volumetric renewal rate of 0.97 ± 0.41 *µ*L/min in the lateral ventricles across the cohort.

As listed in Table 1, statistical evaluation of the fitting quality showed that the goodness-of-fit was significantly higher for the proposed model (*R*^2^ = 0.95 ± 0.05) compared to the conventional framework (*R*^2^ = 0.82 ± 0.06, *p* < 0.001). Additionally, the Akaike Information Criterion (AlC) was significantly reduced from 106.93 ± 2.50 (conventional) to 94.96 ± 5.83 (proposed) (*p* < 0.001).

**Table 1:** Comparison of kinetic parameters and goodness-of-fit metrics between the proposed and conventional models (N=11).

| Parameter | Proposed model<br>(mean ± STD) | Conventional model<br>(mean ± STD) | Relative difference | P-value |
| --- | --- | --- | --- | --- |
| Localized CSF renewal rate, $f'$ ($\times 10^{-3} \text{ s}^{-1}$)<br>or $f$ ($\times 10^{-3} \text{ s}^{-1}$) | $4.37 \pm 1.02$ | $3.81 \pm 0.92$ | + 14.70% | $P < 0.001$ |
| Arterial transit time, $\delta$ (s) | $0.55 \pm 0.08$ | $0.28 \pm 0.07$ | + 96.43% | $P < 0.001$ |
| Goodness-of-fit, $R^2$ | $0.95 \pm 0.05$ | $0.82 \pm 0.06$ | + 15.85% | $P < 0.001$ |
| Akaike information criterion (AIC) | $94.96 \pm 5.83$ | $106.93 \pm 2.50$ | - 11.19% | $P < 0.001$ |

## Discussion

In this study, we developed and validated a dynamically coupled kinetic model to improve ASL-based quantification of CSF dynamics. We built upon the pioneering foundational work[4], which established a robust and essential methodology for evaluating CSF dynamics using the FAIR-ASL sequence. Under the assumption that the lateral ventricular volume remains constant during the ASL acquisition, the volume of fluid influx must exactly equal that of efflux. Guided by this balanced influx-efflux principle, our model requires that physiological clearance (*f*′) be explicitly separated from and dynamically coupled with intrinsic *T*_1_ relaxation (*R*_*eff*_ = *R*_1,*CSF*_ + *f*′), so that the physical magnetic relaxation and biological fluid washout can be distinguished. Furthermore, by introducing a zero-clamp boundary condition (*TI* < *δ*), we eliminated non-physiological negative signal estimations during the tracer arrival phase, ensuring strict adherence to physiological reality.

Beyond refining the theoretical compartment dynamics, our framework resolves a critical sequence-specific discrepancy historically present in FAIR-ASL analyses. Previous evaluations of FAIR datasets applied analytical assumptions suited for other ASL techniques (such as PASL), specifically by fitting a finite bolus duration (*τ*). However, because FAIR utilizes a global inversion pulse, it effectively creates an infinitely long resenvoir of labeled spins (*τ* → ∞) that decays solely via *T*_1_ relaxation. Forcing a finite bolus assumption onto FAIR data creates a fundamental mathematical discrepancy, prompting optimization algorithms to generate artificial curve truncations. By correcting this historical oversight and rigidly anchoring our model to the FAIR sequence physics (*τ* → ∞), we streamlined the model into a two-parameter space (*f*′, *δ*).

The synergy of mass conservation principles and sequence-specific boundary conditions provides a substantially more robust and physiologically accurate representation of CSF dynamics. By eliminating the artificial label truncation, our model reproduces the smooth, continuous washout curves observed in vivo. Consequently, while the conventional constrained framework might underestimate signal exchange dynamics, our sequence-matched formulation recovers these dynamics, yielding higher and more reliable estimates of the localized CSF renewal rate (*f*′) observed in our cohort. This theoretical and practical superiority is strongly supported by significantly improved goodness-of-fit and more favorable Akaike Information Criterion (AIC) metrics. The significantly higher CSF dynamic metric *f*′ values recovered by our model suggest that previous analytical methods may have systematically underestimated CSF dynamics due to artificial mathematical constraints. Accurately capturing the CSF dynamics with a robust, two-parameter model holds significant translational potential, particularly when investigating microscopic CSF dynamics is crucial for understanding conditions like Alzheimer’s disease and impaired glymphatic clearance. Ultimately, rather than oversimplifying the biological process, this framework demonstrates that strict, simultaneous adherence to both fluid kinetics and actual sequence physics is essential for maximizing the reliability of ASL quantification.

The physiological estimates derived from this refined model offer valuable insights into blood-CSF barrier water transport and exchange. Utilizing our mass-conservation model, we estimated a mean localized CSF renewal time in the lateral ventricles (4.01 ± 0.97 minutes) and a corresponding volumetric renewal rate of 0.97 ± 0.41 *µ* L/min across the cohort (*N* = 11). Notably, this volumetric renewal rate is substantially higher than the classical macroscopic CSF production rates reported in mice, which typically range from 0.090 *µ*L/min (measured via direct perfusion in the lateral and third ventricles)[15] to 0.325 *µ* L/min (estimated via indirect perfusion across all ventricles)[16]. However, this apparent discrepancy is fundamentally expected. It is important to emphasize that ASL-derived metrics primarily reflect water transport across the choroid plexus, as well as water exchange across the lateral ventricular wall, rather than the net unidirectional macroscopic bulk flow.

To appropriately interpret our findings, it is essential to distinguish the specific physiological targets of ultra-long TE ASL from other existing CSF imaging modalities. Unlike Phase Contrast MRI (PC-MRI), which exclusively captures macroscopic, pulsatile bulk flow within major conduits and remains fundamentally insensitive to slow, perfusion-like tissue-CSF interactions[17], our ASL framework specifically quantifies molecular-level water transport across the choroid plexus and water exchange across the lateral ventricular wall. Conceptually, this trans-barrier water turnover aligns more closely with the physiological processes probed by isotope tracers such as D_2_O or 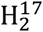 O MRl [18-20]. While those direct tracer methods can precisely profile blood-CSF equilibration time scales or absolute water flux, their broad clinical utility is severely restricted by invasiveness, ionizing radiation, and prohibitive costs. In addition, ASL allows repeated measurements within a single session. Therefore, although the proposed ASL model intrinsically characterizes unidirectional CSF water dynamics rather than net volumetric CSF secretion or bidirectional equilibration, it uniquely fills a critical translational gap. It provides a fully noninvasive, accessible, and parameter-robust methodology to interrogate BCSFB integrity and trans-ependymal water exchange, phenomena that are heavily implicated in neurodegeneration but remain entirely inaccessible to conventional PC-MRI.

Despite these highly promising advancements, several limitations of the current study should be acknowledged. First, the validation of our proposed model was performed on a relatively small open-source cohort (*N* = 11). Future studies incorporating larger, multi-site cohorts are necessary to establish definitive normative ranges for the refined *f*′ and *δ* parameters across different age groups. Second, while the macroscopic constant-volume assumption is mathematically indispensable for enforcing fluid mass conservation, it does not explicitly model microscopic physiological pulsatility (e.g. cardiac- or respiratory-driven transient volume fluctuations). Moreover, this assumption may be less valid under pathological conditions in which lateral ventricular volume changes persistently. However, because the lateral ventricles are surrounded by brain parenchyma, their volume may still reasonably be assumed to remain constant during the measurement period under stable physiological or pathological conditions. The third limitation is the well-mixed assumption for lateral ventricular CSF, which underlies the first-order washout term by assuming that efflux CSF carries the mean compartmental difference magnetization. While this is a reasonable ROI-level simplification, it may break down in pathological conditions associated with ventricular enlargement, compartmentalization, or impaired mixing. Finally, the current *τ* → ∞ formulation is a sequence- and hardware-specific idealization for non-selective FAIR inversion and is most appropriate for the present preclinical implementation. In human FAIR-ASL experiments, limited transmit-field coverage may effectively introduce a finite bolus duration, such that a more general finite-*τ* formulation may be required. Expanding this mass-conservation model to accommodate other labeling strategies (such as PCASL, which indeed has a defined finite *τ*) will require re-integrating the bolus duration while strictly maintaining the influx-efflux balance, this remains an active area for future work. The present model should be interpreted as a sequence-specific, hardware-specific, and compartment-level description of apparent CSF renewal rather than a direct measurement of CSF production. Its assumptions are most appropriate for the current FAIR-ASL acquisition and may require reformulation for finite-bolus labeling sequences.

## Conclusion

In conclusion, by explicitly aligning mathematical modeling with the strict principles of fluid mass conservation and sequence-specific physics, we have overcome significant theoretical limitations in conventional FAIR-ASL analyses of CSF dynamics. This parameter-streamlined, dynamically coupled model eliminates mathematical artifacts and recovers previously constrained CSF dynamics, establishing a highly stable and reliable methodology for non-invasive CSF quantification.

## Supporting information

Theoretical Derivation

Curve fitting

