## Supplementary material for "A dynamically coupled kinetic model for Arterial Spin Labeling: Enhancing the reliability of ventricular cerebrospinal fluid renewal quantification in vivo": Theoretical Derivation

### Derivation of a FAIR-ASL based kinetic model for ventricular blood-to-CSF fluid dynamics under mass conservation

#### I. Modeling rationale and constraints

To quantify blood-to-CSF fluid dynamics in the lateral ventricles using FAIR-ASL, we formulated a compartmental kinetic model under the following constraints:

- The equilibrium magnetization and longitudinal relaxation rate of ventricular CSF are estimated independently from the control signal, such that $M_0$ and $R_{1,CSF}$ are treated as known quantities in the subsequent fitting of the ASL difference signal.
- The transit-delay phase, during which no labeled difference signal has yet reached the ventricular CSF compartment, must be explicitly incorporated.
- Because FAIR-ASL uses a global inversion pulse, the effective labeled bolus duration is assumed to be infinitely long, i.e., $\tau \to \infty$.
- The lateral ventricular volume is assumed to remain constant during the measurement period.
- Under this fixed-volume assumption, the volumetric influx rate equals the volumetric efflux rate.
- Efflux-driven signal loss must be explicitly represented as a separate physiological process and should not be phenomenologically absorbed into a single relaxation constant. Under these assumptions, the ASL difference signal is modeled as a tracer-like difference-magnetization quantity within the ventricular CSF compartment.

#### II. Fixed-volume mass conservation

Let $V_{\mathrm{LV}}$ denote the combined volume of the lateral ventricles, which is assumed to remain constant over the acquisition window. Let $Q$ denote the volumetric influx rate of fluid entering the ventricle. Under the fixed-volume assumption, influx must equal efflux, such that

$$Q_{\mathrm{in}} = Q_{\mathrm{out}} = Q \tag{1}$$

We define the localized CSF renewal rate $f'$ as the fractional replacement:

$$f' = \frac{Q}{V_{\mathrm{LV}}} \tag{2}$$

Thus, the volumetric influx rate can be written as

$$Q = f^{'} V_{\mathrm{LV}} \tag{3}$$

Accordingly, under the assumptions of constant ventricular volume and balanced influx-efflux, the efflux-driven replacement rate per unit compartment volume is also given by $f'$. This allows the washout term to be expressed directly as a first-order loss process proportional to the existing ventricular difference magnetization.

**III. Parameter definitions**

The model variables and parameters are defined as follows:

- $\Delta M(t)$: ASL difference signal in the lateral ventricular CSF compartment, defined here as the label-control magnetization difference $\Delta M = M_{label} - M_{control}$ attributable to inflowing unlabeled water.
- $f'$: localized CSF renewal rate, defined as the fractional replacement rate of ventricular CSF water by blood-derived water per unit lateral ventricular volume; that is, the rate at which pre-existing ventricular CSF water is replaced by newly arriving blood-derived water.
- $M_0$: equilibrium magnetization of ventricular CSF, estimated independently from the control signal.
- $R_{1,CSF}$: longitudinal relaxation rate of ventricular CSF, estimated independently from the control signal.
- $\delta$: transit delay for blood-derived water to reach the ventricular CSF compartment.
- $\varphi$: blood-to-CSF partition coefficient.
- $\alpha$: labeling efficiency.
- $R_{1a}$: longitudinal relaxation rate of arterial blood.

**IV. Governing kinetic equation**

We model the ventricular ASL difference signal $\Delta M(t)$ as a compartmental tracer-like quantity, rather than as the absolute longitudinal magnetization in either the label or control image alone. Under the assumptions of fixed ventricular volume and balanced influx-efflux, the temporal evolution of $\Delta M(t)$ is governed by three components: (i) influx of FAIR-ASL generated difference magnetization after a transit delay $\delta$, (ii) intrinsic longitudinal relaxation within the CSF compartment at rate $R_{1,CSF}$, and (iii) physiological washout due to fluid efflux at rate $f'$. Accordingly, the governing equation is

$$\frac{d\Delta M(t)}{dt} = \mathrm{Influx}(t) - R_{1,CSF}\,\Delta M(t) - f'\,\Delta M(t) \tag{4}$$

This can be rearranged as

$$\frac{d\Delta M(t)}{dt} = \mathrm{Influx}(t) - \left(R_{1,CSF} + f'\right)\,\Delta M(t) \tag{5}$$

Although $R_{1,CSF}$ and $f'$ appear additively in the governing equation, they represent distinct physical processes: intrinsic $T_1$ relaxation of CSF magnetization and physiological clearance by fluid efflux, respectively. They should therefore remain conceptually separate rather than being collapsed into a single fitted apparent decay constant.

##### 4.1 Expression of the influx term

When $t < \delta$, the unlabeled blood water has not yet reached the lateral ventricles, so the influx term is zero. For $t \geq \delta$, the influx is continuously supplied and is not truncated by a finite bolus under the assumption $\tau \to \infty$. The inversion-induced difference magnetization in arterial blood decays according to arterial $T_{1,a}$, or equivalently at rate $R_{1a}$, so the difference magnetization input is written as,

$$J_{in}(t) = \frac{2M_0 f'\alpha}{\varphi} e^{-R_{1a}t} H(t-\delta) \tag{6}$$

where $H(t-\delta)$ is the Heaviside step function. Note that the parameter $f'(\mathrm{s}^{-1})$ represents the localized CSF renewal rate, defined as the fractional replacement rate of ventricular CSF water by blood-derived water per unit lateral ventricular volume; that is, the rate at which pre-existing ventricular CSF water is replaced by newly arriving blood-derived water. This form is consistent with the input amplitude in the original Evans model. The factor of 2 reflects the inversion difference between control and label conditions in the FAIR-ASL sequence.

The influx term represents the transport of difference magnetization into the ventricular CSF compartment, with amplitude determined by the FAIR-ASL preparation and temporal decay governed by arterial $T_1$.

##### 4.2 $T_1$ decay term

The longitudinal relaxation rate of ventricular CSF, $R_{1,CSF}$, is estimated independently from the control signal and is held fixed during the subsequent fitting of the difference signal. The corresponding relaxation term is:

$$-R_{1,CSF}\Delta M(t) \tag{7}$$

This separation is a modeling assumption introduced to preserve the physiological interpretability of $f'$.

##### 4.3 Efflux term

The loss of difference magnetization due to fluid efflux is proportional to the difference magnetization currently present in the ventricular compartment and is therefore written as

$$-f'\Delta M(t) \tag{8}$$

This first-order washout form assumes that the ventricular CSF compartment is well mixed, such that the effluent carries the same mean difference magnetization concentration as the compartment itself.

##### 4.4 Final differential equation

Substituting (6)–(8) into (4), we get:

$$\frac{d\Delta M(t)}{dt} = \frac{2M_0 f'\alpha}{\varphi} e^{-R_{1a}t} H(t-\delta) - R_{1,CSF}\Delta M(t) - f'\Delta M(t) \tag{9}$$

which rearranges to:

$$\frac{d\Delta M(t)}{dt} + \left(R_{1,CSF} + f'\right)\Delta M(t) = \frac{2M_0 f'\alpha}{\varphi} e^{-R_{1a}t} H(t-\delta) \tag{10}$$

Note that although $R_{1,CSF}$ and $f'$ appear additively in the governing differential equation, they represent fundamentally distinct processes and should not be interpreted as a single undifferentiated apparent decay constant.

#### V. Piecewise solution process

The solution is first derived as a function of continuous time $t$ and is then evaluated at $t = TI$.

##### 5.1 Phase 1: $TI < \delta$

When $t < \delta, H(t-\delta) = 0$, so the input term is 0. Assuming the initial condition $\Delta M(0) = 0$, we obtain:

$$\Delta M(TI) = 0, \quad TI < \delta \tag{11}$$

##### 5.2 Phase 2: $TI \geq \delta$

When $t \geq \delta$, the equation becomes:

$$\frac{d\Delta M(t)}{dt} + \left(R_{1,CSF} + f'\right)\Delta M(t) = \frac{2M_0 f'\alpha}{\varphi} e^{-R_{1a}t} \tag{12}$$

with the boundary condition:

$$\Delta M(\delta) = 0$$

##### 5.3 Solving with the integrating factor method

The integrating factor is chosen as:

$$\mu(t) = e^{(R_{1,CSF}+f')t} \tag{13}$$

Multiplying both sides by $\mu(t)$:

$$e^{(R_{1,CSF}+f')t}\frac{d\Delta M}{dt} + \left(R_{1,CSF} + f'\right)e^{(R_{1,CSF}+f')t}\Delta M = \frac{2M_0 f'\alpha}{\varphi} e^{(R_{1,CSF}+f'-R_{1a})t} \tag{14}$$

The left-hand side can be written as an exact derivative:

$$\frac{d}{dt}\left[\Delta M(t)e^{(R_{1,CSF}+f')t}\right] = \frac{2M_0 f'\alpha}{\varphi} e^{(R_{1,CSF}+f'-R_{1a})t} \tag{15}$$

Integrating from $\delta$ to $TI$:

$$\Delta M(TI)e^{(R_{1,CSF}+f')TI} - \Delta M(\delta)e^{(R_{1,CSF}+f')\delta} = \frac{2M_0 f'\alpha}{\varphi}\int_{\delta}^{TI} e^{(R_{1,CSF}+f'-R_{1a})t}\,dt \qquad (16)$$

Since $\Delta M(\delta) = 0$, we have:

$$\Delta M(TI)e^{(R_{1,CSF}+f')TI} = \frac{2M_0 f'\alpha}{\varphi}\int_{\delta}^{TI} e^{(R_{1,CSF}+f'-R_{1a})t}\,dt \qquad (17)$$

5.4 Calculating the integral

Let $\Lambda = R_{1,CSF} + f' - R_{1a}$, then:

$$\int_{\delta}^{TI} e^{\Lambda t}\,dt = \frac{e^{\Lambda TI} - e^{\Lambda\delta}}{\Lambda} \qquad (18)$$

Substituting this back into (17) and expanding:

$$\Delta M(TI) = \frac{2M_0 f'\alpha}{\varphi} e^{-(R_{1,CSF}+f')TI} \cdot \frac{e^{(R_{1,CSF}+f'-R_{1a})TI} - e^{(R_{1,CSF}+f'-R_{1a})\delta}}{R_{1,CSF}+f'-R_{1a}}, \quad TI \geq \delta \qquad (19)$$

**VI. Further simplification of the formula**

Because $e^{-(R_{1,CSF}+f')TI} \cdot e^{(R_{1,CSF}+f'-R_{1a})TI} = e^{-R_{1a}TI}$, we can simplify and rearrange the equation into a more physically meaningful form:

$$\Delta M(TI) = \frac{2M_0 f'\alpha}{\varphi} \cdot \frac{e^{-R_{1a}TI} - e^{-R_{1a}\delta} e^{-(R_{1,CSF}+f')(TI-\delta)}}{R_{1,CSF}+f'-R_{1a}}, \quad TI \geq \delta \qquad (20)$$

**VII. Final piecewise model**

Combining the conditions for $TI < \delta$ and $TI \geq \delta$ the final complete model is:

$$\Delta M(TI) = \begin{cases} 0, & TI < \delta \\ \frac{2M_0 f'\alpha}{\varphi} \cdot \frac{e^{-R_{1a}TI} - e^{-R_{1a}\delta} e^{-(R_{1,CSF}+f')(TI-\delta)}}{R_{1,CSF}+f'-R_{1a}}, & TI \geq \delta \end{cases} \qquad (21)$$

**VIII. Normalized form**

If the normalized signal $\Delta M/M_0$, is fitted, the model can be written as:

$$\frac{\Delta M(TI)}{M_0} = \begin{cases} 0, & TI < \delta \\ \frac{2f'\alpha}{\varphi} \cdot \frac{e^{-R_{1a}TI} - e^{-R_{1a}\delta} e^{-(R_{1,CSF}+f')(TI-\delta)}}{R_{1,CSF}+f'-R_{1a}}, & TI \geq \delta \end{cases} \qquad (22)$$

If you use the $M_0^{corr}$ normalization mentioned in Evans' paper, simply replace the $M_0$ in the denominator with $M_0^{corr}$.

**IX. Interpretation**

A key feature of this formulation is that the apparent decay of the ventricular ASL difference signal is not treated as a purely empirical fitting constant. Instead, the model explicitly separates:

- **intrinsic CSF relaxation**, governed by $R_{1,\mathrm{CSF}}$, from
- **physiological renewal/washout**, governed by $f'$.

Although these two processes appear additively in the governing differential equation, they reflect fundamentally different mechanisms. This distinction is essential for preserving the physiological interpretability of the fitted renewal rate. In addition, the explicitly imposing the pre-arrival condition $\Delta M(t) = 0$ for $t < \delta$ prevents non-physical negative signal estimates at very short inversion times, while the FAIR-ASL specific assumption $\tau \to \infty$ avoids artificial truncation associated with finite-bolus models that are not sequence-matched to FAIR-ASL physics.

This formulation is a lumped-compartment model valid under the assumptions of fixed ventricular volume, balanced influx-efflux, a well-mixed ventricular ROI, and FAIR-ASL consistent effectively infinite bolus duration. It does not explicitly model spatial transport, finite-bolus truncation, or separate exchange pathways across distinct ventricular interfaces.
