## Supplementary material for "A dynamically coupled kinetic model for Arterial Spin Labeling: Enhancing the reliability of ventricular cerebrospinal fluid renewal quantification in vivo": Curve fitting

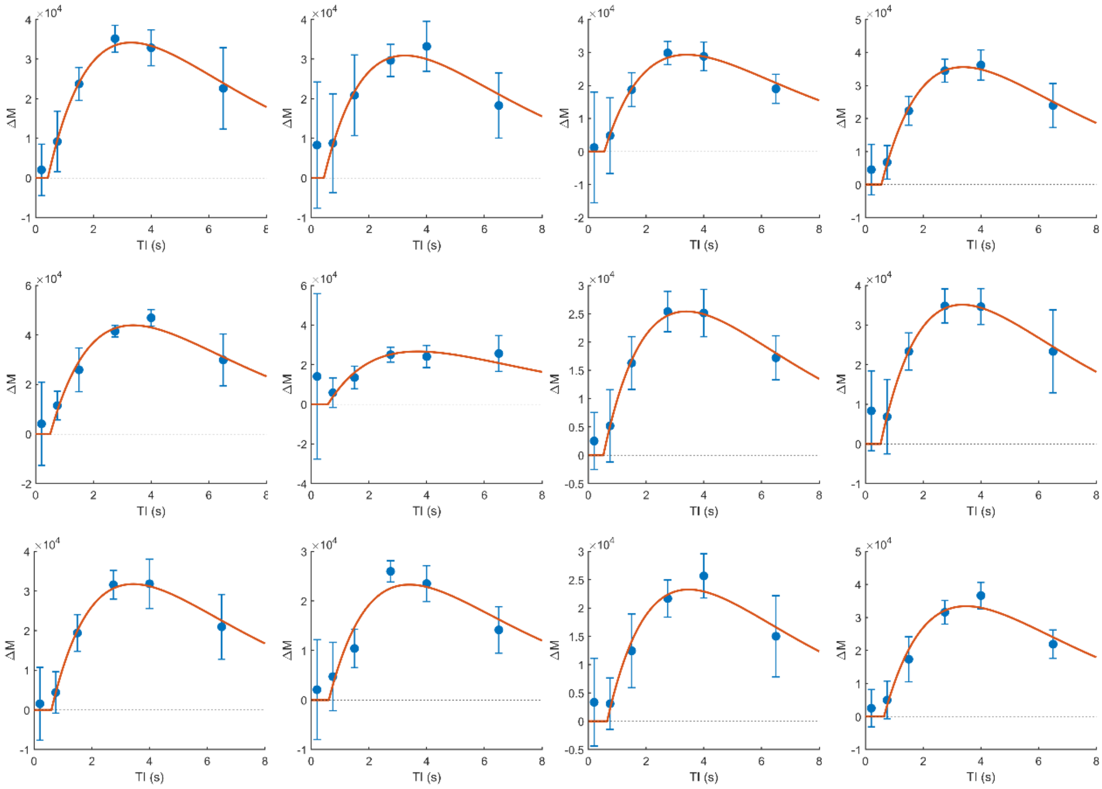


Figure S1. All fitted curves using the proposed two-parameter model. Following data quality review, the subject in row 2, column 2 was excluded due to an artifact at the first T1, yielding a final cohort of N = 11 for quantitative analysis.
